# Therapeutic potential of glabridin and gymnemic acid alleviates eye choroidal thickness and neovascularization in diabetic model rats

**DOI:** 10.1101/2024.02.08.579467

**Authors:** Manaras Komolkriengkrai, Udomlak Matsathit, Wipapan Khimmaktong

## Abstract

Small blood vessels in the eyes are more susceptible to injury, which can lead to complications. However, since diabetic retinopathy is often a serious clinical condition, most of this study focuses on the vascular system of the choroid. As part of this study, we looked at how gymnemic acid (from *Gymnema sylvestre*) and glabridin (from *Glycyrrhiza glabra*, or licorice) might help diabetic rats’ choroid thickness and blood vessels. The aim of this study was to explore the effects of glabridin and gymnemic acid on the structural changes of the choroidal layer as well as the expression of VEGF and CD31 in diabetic rats. The animals were separated into five groups: the control group (Control), the diabetic group (DM), the diabetic rats treated with glabridin 40 mg/kg body weight (DM+GB), the diabetic rats treated with gymnemic acid 400 mg/kg body weight (DM+GM), and the diabetic rats treated with glyburide 4 mg/kg body weight (DM+GR). There was an increase in the thickness of both the choroid layer and the wall of the arteries in the diabetic group. A decrease in vascularity and choroidal neovascularization was found in DM rats. After eight weeks of experimentation, the choroidal thickness increased and the walls of choroid arteries, as well as the diameter of the lumen of choroid arteries in the DM+GB, DM+GM, and DM+GR groups, were wider. The expression of VEGF and CD31 was lower compared to the DM group. According to the findings of the current research, glabridin and gymnemic acid can reduce the damage to the choroid, which is a factor that can sometimes result in vision loss.

## Introduction

Diabetes issues can eventually result in blindness and give rise to the serious condition known as diabetic retinopathy (DR). High blood sugar makes the vascular wall less effective because it kills intramuscular pericytes and makes the basement membrane thicker (1). This happens when microvascular retinal changes occur. The small blood vessels in the eyes are particularly vulnerable to poor blood glucose control, which can cause damage (2). Studies on humans and animals have mostly examined the retinal vasculature rather than the choroid vasculature, since diabetic retinopathy is typically a serious clinical issue.

The choroid, which is located outside the retinal pigment epithelium (RPE) and supplies the outer retina with nutrients, normally supplies the inner retina through the retinal vasculature. Therefore, it is possible that the thickness of the choroid may indirectly reflect the metabolic condition of the retina and the circulation of the choroidal blood. Several retinal illnesses, including glaucoma, have been linked to abnormalities in the thickness of the choroidal layer. Therefore, the nonperfusion of choroidal capillaries or the choriocapillaris could result in functional vision loss (3). In addition to supplying the outer retina with nutrients, the choroid is responsible for regulating both the temperature and the volume of the eye. The choroidal circulation is a high-flow system that has a relatively low oxygen content, even though it is responsible for 85% of the total blood flow in the eye. The posterior choroid and the peripapillary region receive their blood supply from the short PCA, while the anterior sections of the choroid receive their blood supply from the long PCAs as well as the anterior ciliary artery. The ophthalmic artery gives off a branch called the anterior ciliary artery, which supplies blood to the iris as well as the anterior choriocapillaris.

Even though diabetic retinopathy is well understood and frequently used as a diagnostic tool to track the advancement of the illness, a routine clinical ophthalmology evaluation of the choroid is quite uncommon (4). According to new research, the diabetic choroid may experience comparable phenomena (5). Also, diabetic retinopathy has been seen in alloxan and streptozotocin (STZ) diabetes models (6). The rats and monkeys that develop diabetes on their own show choroidal vascular leakage and capillary dropout and display evidence of choroidal neovascularization. The thickening of the basement membrane coined the term diabetic choroidopathy (7). It is possible that choroidal vasculopathy that results from diabetes plays a significant role in the development of diabetic retinopathy. Diabetic choroidopathy may also be the cause of the inexplicable loss of visual function that can happen to diabetic subjects who do not have retinopathy (5).

In experimental models like type 1 diabetes, streptozotocin (STZ) is a commonly used drug to generate insulin-dependent diabetic mellitus. STZ causes potentially dangerous systemic microvascular changes and consequences. It also has toxic effects on islet beta cells. The vascular corrosion casting technique combined with scanning electron microscopy (SEM) has become an important basic tool for learning about how organ microvasculation works. This will help future physiological and pathological studies. The three-dimensional architecture of the vascular bed and network can be identified using this conventional method (8). This technique produces a readily comprehensible, clear, and identifiable three-dimensional picture of the entire choroid as well as any enlarged localized lesions, all while revealing the vascular architecture.

At present, modern methods of treating diabetic eye disease have many effects on patients. In addition, as the cost of treatment is quite expensive and can result in various side effects, there a currently much research on herbs that have the power to cure diabetes. Gymnemic acid is an important substance in *Gymnema sylvestre*, which is an herb used to treat diabetes (9). It has the effect of increasing the efficiency of β-cell function in the pan-creas and slowing down the absorption of glucose in the small intestine. This results in an increase in the amount of insulin in the body and can reduce blood sugar levels in diabetic rats (10). Currently, there are research studies on diabetes that affect various organs in the body. In this paper, we studied the action of gymnemic acid as an alternative option for treating diabetic conditions. Within the scope of this research project, the effects of gymnemic acid derived from *Gymnema sylvestre* on microvascularity and structural issues within the choroids of rats that had been given STZ to induce diabetes were investigated.

Glabridin is a polyphenolic flavonoid that is also recognized as an active component of the Glycyrrhiza glabra, which is also known as the licorice plant. Streptozotocin produced diabetic rats have shown that it lowers blood glucose levels (11). Addition-ally, it has demonstrated evidence of having an antioxidant effect in the kidneys of diabetic’s mice through an increase in superoxide dismutase (SOD) and a decrease in malondialdehyde (MDA) content (12). The use of glabridin in diabetic rats resulted in a reduction in serum glucose as well as hepatic collagen type I, which caused the livers of the diabetic rats to return to their normal structures (11). Because of this, glabridin’s ability to lower blood sugar and fight inflammation and scar tissue may make it an important component of future drug development plans for diabetes. For that reason, in this study, we used a microscope to examine how gymnemic acid and glabridin affect the blood vessels in the choroidal layer of diabetic rats’ eyes and compare the levels of VEGF and CD31 proteins after gymnemic acid treatment. These two proteins cause angiogenesis, a process that can result in eye damage and diabetic retinopathy. Therefore, it is expected that this research project will yield multiple benefits and be a guideline for using herbs to prevent diabetes related eye complications from causing vision loss.

## Materials and Methods

### Animals and the methodology of the study

In this investigation, we used male Wistar rats that were eight weeks old and weighed between 200 and 250 g. At the Southern Laboratory Animal Facility at Prince of Songkla University, a total of fifty rats were housed in an environmentally controlled laboratory environment. The conditions included a humidity level of 50 + 10%, a temperature of 25 + 2°C, and a light/dark cycle of 12 hours on/12 hours off. The Wistar rats had unrestricted access to both the food and the water from the tap. The Prince of Songkla University Animal Ethics Committee gave its stamp of approval to every experimental technique that was carried out utilized (MHCSI 6800.11/236, Ref.3/2020).

### Initiation of diabetes in animals

One week after becoming accustomed to their new environment, the rats were divided at random into five groups (each with ten members). In the control group, the rats received a standard rat diet. The standard diet was given to the rats in the diabetic group, which was denoted by the acronym STZ (streptozotocin). The diabetic rats in the DM+GB group were given a standard diet and were also given glabridin from licorice 4 mg/kg BW (purified > 98% via HPLC analysis performed by Shaanxi Langrun Biotechnology Co., Ltd., Xian, China) in 0.5 ml of 0.5% Tween 80 solution as a supplement. The diabetic rats in the DM+GM group were given a standard diet and were also given gymnemic acid from *Gymnema sylvestre* 400 mg/kg BW (purified > 75% via HPLC analysis) in 0.5 ml of 0.5% Tween 80 solution as a supplement. The diabetic rats in the DM+GR group were given a standard diet and were also given glyburide (Sigma-Aldrich; Merk KGaA) at 4 mg/kg in 0.5 ml of 0.5% Tween 80 solution as a supplement. To induce hyperglycemia, a single dosage of streptozotocin (STZ), i.e., 60 mg/kg (Sigma-Aldrich; Merk KgA, Germany), that was dissolved in 0.1 mol/l in citrate buffer was administered intraperitoneally (I.P.) to all rats, except for those in the control group of rats. Citric buffer was the only substance that was administered to the control rats. Blood glucose levels were tested using a glucometer manufactured by Roche Diagnostics GmbH in Mannheim, Germany, 3 days after an injection of STZ. The rats that had a blood glucose level higher than 250 mg/dl were categorized as diabetic rats. The level of glucose in the blood was measured once a week. Eight weeks after receiving therapy, the rats were sacrificed under anesthesia using an excessive dose of sodium pentobarbital (200 mg/kg; intraperitoneal injection), and a blood sample was taken from their hearts using a cardiac puncture. After dissection, the eyes were placed in 10% formalin buffer for preservation.

### Preparation of histological slides for Masson’s trichrome staining procedure

As a part of the preparation for the histological investigation, the eye tissues of all the groups were dissected and were then promptly fixed in 10% formalin. This was carried out to assess the histological alterations and measure the lumen diameter of the choroidal arteries. These were subjected to a graded sequence of ethanol that progressed through 70, 80, 90, 95, and 100% for one hour each, with two changes in between. Before filtering, three changes of xylene lasting thirty minutes each were employed as a cleaning reagent. The tissue was then fixed in paraffin, cut into sections that were five micrometers thick, and stained with Masson’s trichrome (Sigma-Aldrich; Merck KGaA). An Olympus light microscope, model BX-50, manufactured in Japan by Olympus, was used to inspect, and photograph each segment. The Olympus cellSens software was used to obtain a measurement of the lumen diameter of the arteries.

### The immunofluorescence method was utilized to investigate the VEGF and CD31 concentrations

An immunohistochemical approach was used to examine the levels of VEGF and CD31 expression in the choroid layer of the eye. Briefly, each slide was deparaffinized in xylene, hydrated in gradient ethanol, permeabilized in PBS buffer, and then blocked with serum in PBS. Finally, the slides were washed in PBS. At a temperature of 4 °C for one night, the slides were treated with rabbit anti-VEGF and mouse anti-CD31 antibodies from Abcam, Cambridge, UK, diluted at 1:200 in blocking serum. The sections were washed three times with PBS before being placed in blocking solution with a fluorescein horse anti-mouse IgG (H+L) antibody and a Texas red goat anti-rabbit IgG (H+L) antibody (1:200; Vector Laboratories, Inc.). They were left there for two hours at room temperature to find VEGF and CD31. A fluorescent microscope (model BX-50; manufacturer: Olympus, Japan) was used to evaluate the images. Within each specimen, five photos at 600-fold magnification in standard fields of 810,965 μm2 were chosen at random for examination. Image J software from the National Institutes of Health (NIH) was used to quantify the chemiluminescence intensity. The optical density (OD) of each sample was standardized based on the photo at 600-fold area. The photos were selected after being grayscaled with 8 bits and then transformed. To obtain accurate measurements of the area, integrated density, and background intensity, the optical density (OD) calculation followed Fu et al.’s (2018) instructions, as shown in the choroidal layer (13).

### An investigation of the choroid vascular corrosion cast process produces a replica of vascular beds using a scanning electron microscopy (SEM)

The left ventricle of each group of rats was perfused with a 0.9% sodium chloride solution to flush the blood out of the blood vessels. After that, a combination of plastic called PU4ii casting resin, which is based on polyurethane (vasQtec), was injected into the blood circulation of rats. To ensure that the plastic was properly polymerized, the animal that had plastic injected into it was first left at room temperature for 30 minutes, and then it was submerged in hot water for 3 hours. After the polymerization process was complete, the eye was extracted, and the tissue was then corroded in a 10% KOH solution at room temperature for 30–40 days. The vascular cast for the eye was given a rinsing under a tap with a gentle stream of water and then washed in multiple changes of distilled water to remove any leftover tissues. After that, it was allowed to air dry at ambient temperature, and then it was mounted to a metal stub using double glue tape and carbon paint before it was coated with gold using a sputtering apparatus. Finally, an SEM (JEOL JSM-5400) with a 10 KV accelerating voltage was used examine the choroid area of the eye cast. A piece of software called SemAfore was used to determine the diameter of the replica vascular bed in the choroid.

### Examination of data based on statistics

The findings were summarized using the mean value together with the standard error of the mean (SEM). The study of statistics was carried out with the use of ANOVA, and then the Bonferroni posttest was used. It was determined that statistical significance was reached when the value of p was lower than 0.05.

## Results

### The influence of glabridin and gymnemic acid on the levels of glucose and HbA1c in the blood

The blood sugar level was measured every week of the experiment for 8 weeks and HbA1c was measured after 8 weeks experiment. It was found that the blood sugar level (Table 1) and HbA1c (Table 2) of the rats in the diabetes group (DM) increased significantly. Statistically significant (p < 0.001 and p < 0.0001, respectively) when compared to the rats in the control group. In the 8th week of the experiment, the rats in the diabetes group (DM) had blood sugar levels equal to 353.83±35.16 mg/dl, while the control group had a blood sugar level of 74.67±3.31 mg/dl (Table 1).

**Table 1.** A comparison of blood glucose levels in different groups. (mg/dl).

| Wk. | C | DM | DM+GB | DM+GM | DM+GR |
| --- | --- | --- | --- | --- | --- |
|  | (mg/dl) | (mg/dl) | (mg/dl) | (mg/dl) | (mg/dl) |
| 1 | $70.5 \pm 4.92$ | $360.50 \pm 32.33^*$ | $314.83 \pm 30.71^*$ | $362.89 \pm 49.29^*$ | $297.67 \pm 20.69^{**}$ |
| 2 | $72.00 \pm 5.49$ | $290.50 \pm 34.44^*$ | $311.67 \pm 49.73^*$ | $269.22 \pm 35.60^*$ | $241.00 \pm 36.43^{\#}$ |
| 3 | $79.17 \pm 3.25$ | $351.17 \pm 43.25^*$ | $305.33 \pm 52.59^*$ | $188.00 \pm 41.51^*$ | $176.50 \pm 38.65^{\#}$ |
| 4 | $73.33 \pm 3.57$ | $253.67 \pm 30.75^*$ | $170.67 \pm 41.11$ | $240.89 \pm 48.70^*$ | $187.50 \pm 45.61$ |
| 5 | $72.00 \pm 3.98$ | $301.50 \pm 55.41^*$ | $137.00 \pm 40.87^{##}$ | $289.78 \pm 50.06^{##}$ | $192.50 \pm 54.94^{**}$ |
| 6 | $73.58 \pm 3.55$ | $351.00 \pm 34.82^*$ | $69.50 \pm 13.69^{\#}$ | $249.78 \pm 49.68^{##}$ | $149.00 \pm 47.48^{\#}$ |
| 7 | $73.00 \pm 4.25$ | $457.00 \pm 20.99^*$ | $158.33 \pm 37.64^{\#}$ | $259.00 \pm 45.53^{##}$ | $165.83 \pm 49.04^{\#}$ |
| 8 | $74.67 \pm 3.31$ | $353.83 \pm 35.16^*$ | $174.50 \pm 30.99^{\#}$ | $213.44 \pm 40.73^{##}$ | $160.67 \pm 52.46^{\#}$ |

\* $p < 0.001$ compared with the control group.
$\#p < 0.001$ , $##p < 0.01$ compared with the DM group.

**Table 2.**
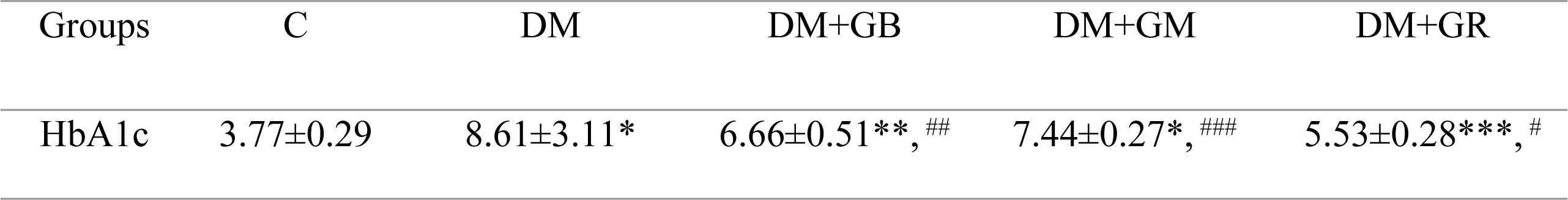
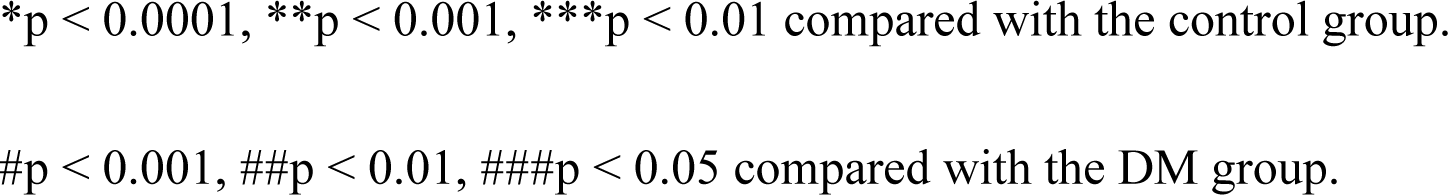
A comparison of HbA1c in different groups.

| Groups | C | DM | DM+GB | DM+GM | DM+GR |
| --- | --- | --- | --- | --- | --- |
| HbA1c | $3.77 \pm 0.29$ | $8.61 \pm 3.11^*$ | $6.66 \pm 0.51^{**}, ##$ | $7.44 \pm 0.27^*, ###$ | $5.53 \pm 0.28^{***}, \#$ |

\*p < 0.0001, \*\*p < 0.001, \*\*\*p < 0.01 compared with the control group.
#p < 0.001, ##p < 0.01, ###p < 0.05 compared with the DM group.

### The effect that glabridin and gymnemic acid have on the total body weight of rats

After the rats induced with diabetes were treated with glabridin (DM+GB), gymnemic acid (DM+GM), and glibenclamide (DM+GR), it was found that the animals in all three groups of these groups displayed a statistically significant decrease in blood sugar levels (p < 0.001, p < 0.01, p < 0.001, respectively) when compared to the rats in the diabetes group (DM), with the mean sugar levels of the rats being equal to 174.50 + 30.99, 213.44 ± 40.73, and 160.67 + 52.46 mg/dl, respectively (Table 3).

**Table 3.** A comparison of the body weights in different groups (grams)

| Wk. | C (g) | DM (g) | DM+GB (g) | DM+GM (g) | DM+GR (g) |
| --- | --- | --- | --- | --- | --- |
| 1 | $256.66 \pm 5.25$ | $178.83 \pm 3.56^*$ | $173.33 \pm 4.71^*$ | $240.83 \pm 4.24^{*****}$ | $197.66 \pm 8.24^*$ |
| 2 | $286.50 \pm 6.17$ | $187.33 \pm 5.52^*$ | $181.16 \pm 5.93^*$ | $216.38 \pm 10.03^{***}$ | $221.33 \pm 14.33^{****}$ |
| 3 | 305.66±8.55 | 178.83±7.60* | 181.00±8.09* | 232.33±8.13*** | 215.50±19.72**** |
| 4 | 325.50±8.17 | 167.50±10.45* | 179.16±9.15* | 231.66±9.19** | 211.33±11.81** |
| 5 | 342.66±9.02 | 171.83±12.38# | 166.50±13.52 | 263.33±15.15## | 212.16±10.28#### |
| 6 | 352.50±8.01 | 173.50±11.01# | 179.16±13.28 | 267.50±15.31## | 209.33±7.85#### |
| 7 | 370.50±8.66 | 159.50±13.52# | 175.50±13.04 | 275.83±16.86## | 199.00±8.03#### |
| 8 | 371.00±9.27 | 182.00±13.82# | 191.00±14.61 | 286.66±18.28### | 222.83±6.25#### |

\*p < 0.000001, \*\* p < 0.0001, \*\*\*p < 0.001, \*\*\*\*p < 0.01, \*\*\*\*\*p < 0.1 compared with the control group.
#p < 0.000001, ##p < 0.001, ###p < 0.01, ####p < 0.1 compared with the DM group.

In the study of the body weight of laboratory rats, the rats were weighed every week of the experiment for 8 weeks. It was found that in the 8th week of the experiment, the body weight of the rats in the diabetes group (DM) decreased significantly (p < 0.001) when compared to the mice in the control group. In the 8th week of the experiment, the mice in the diabetes group (DM) had an average body weight of 165.33+6.56 grams, while the control group had an average body weight of 354.67 + 17.91 grams. After being treated with glabridin (DM+GB), gymnemic acid (DM+GM), and glibenclamide (DM+GR) in rats induced with diabetes, it was found that in all three groups of rats there was an increase in mean body weight compared to the rats in the diabetes group (DM), with the mean body weight of the rats being 185.33 + 6.01, 171.33 + 3.38, and 186.33 + 4.37 grams (Table 3).

### Histological changes in the choroidal blood vessels at the light microscope level

The choroid layer is characterized by loose connective tissue and a high number of blood vessels that serve as its source of nourishment. Within this layer, an intricate network of blood vessels may be found. At the end of the 8-week experiment, eye tissues were stained with Masson’s trichrome (Fig 1). It was found that there was no pathology in the choroidal tissue or blood vessels in the control group (Fig 1). The average thickness of the choroid layer in the control group was 16.56 ± 0.56 µm (Fig 2). It was determined through the process of measuring the dimensions of the choroid artery that the average thickness of the arterial wall was 9.48 ± 1.13 µm, while the lumen width was measured as being 50.47 ± 11.37 µm. High blood sugar levels caused diabetes in the DM group of rats, which had the pathological characteristics of the choroidal tissue and blood vessels (Fig 1). The walls of the blood vessels showed that collagen fibers were building up all over the choroidal and scleral tissues. The thickness of the choroid layer and wall of choroidal arteries increased, exhibiting an increased thickness of collagen fibers which caused the lumen diameter to narrow. The average thickness of the choroid layer in DM rats was 27.22 ± 0.87 µm (Fig 2). The mean choroidal artery wall thickness increased by 16.35 ± 5.01 µm compared to the control group. The mean vascular lumen width was 39.12 ± 9.27 µm, as shown in Table 4.

**Fig 1.**
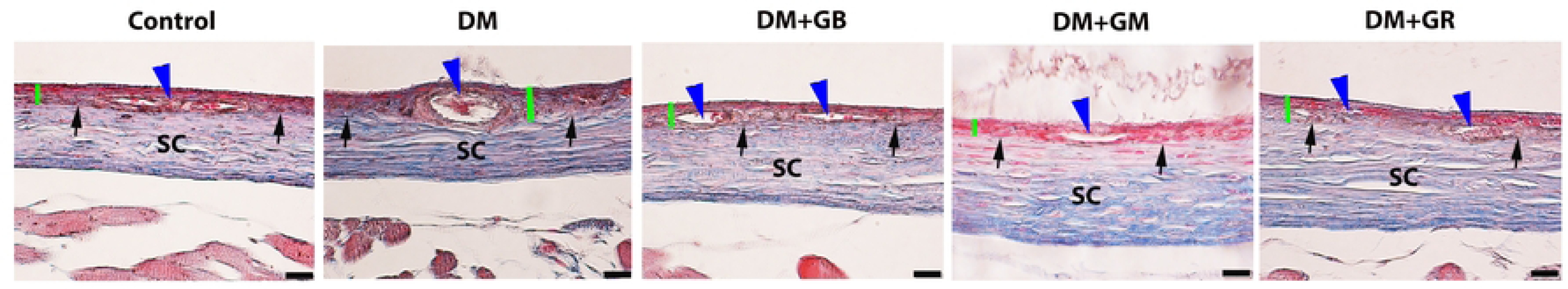
Grabridin and gymnemic acid have ability to reverse histopathological changes in the choroid thickness of diabetic rats. Representative images of the choroidal and scleral (SC) layers of the eye. The choroid and sclera can be separated by choroidal-scleral junctions (black arrows) stained with Masson’s trichrome in all five groups of rats. The thickness of the choroidal layer in each group is presented with the green line. The choroidal layer is composed of choroid arteries (blue arrowheads). Changes in the histological characteristics of the choroidal layer were observed in the DM group, involving the thickness of the choroidal layer and the choroidal arterial wall. Scale bar = 20 µm.

**Fig 2.**
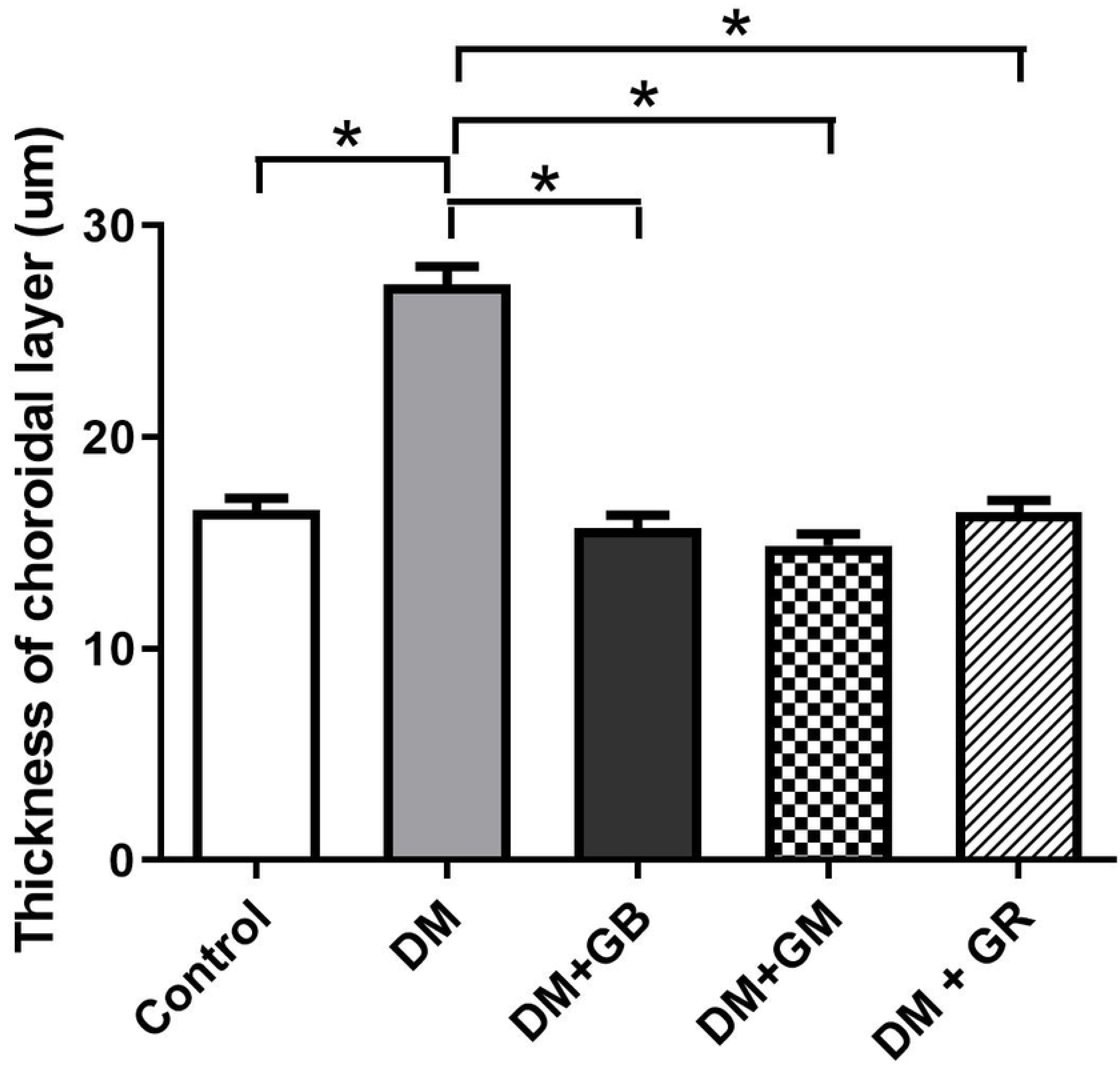
The effect of glabridin and gymnemic acid on the thickness of the choroidal layer in five groups of rats at 8 weeks. A representative graph presents the significant increases in the thickness of the choroidal layer in the DM group compared with the control rats (*p<0.001). After supplementation with glabridin, gymnemic acid, and glyburide, the thickness of choroidal layer decreased in the DM+GB, DM+GM, and DM+GR groups compared with the DM rats (#p<0.001). Values are means + SE.

**Table 4.** The mean wall thickness, and lumen diameter of the choroid arteries (CRA) in all five groups of rats: control group, DM group, DM+GB group., DM+GM group, and DM+GR group at 8 weeks.

| Groups | Wall thickness of CRA ($\mu\text{m}$) | Diameter of lumen of CRA ($\mu\text{m}$) |
| --- | --- | --- |
| Control | $9.48 \pm 1.13^*$ | $50.47 \pm 11.37^*$ |
| DM | $16.35 \pm 5.01$ | $39.12 \pm 9.27$ |
| DM+GB | $10.23 \pm 1.11^*$ | $52.52 \pm 11.10^*$ |
| DM+GM | $10.41 \pm 1.44^*$ | $53.71 \pm 7.12^*$ |
| DM+GR | $9.80 \pm 1.78^*$ | $51.58 \pm 8.11^*$ |

\*p-value < 0.05 compared with the DM group.

On the other hand, when glabridin, gymnemic acid, and glyburide were given to the DM+GB group, DM+GM group, and DM+GR group (Fig 1), these rats displayed thinner choroidal layers than the DM rats (Fig 7). It was found that blood vessels were reduced in choroid artery wall pathology and restored to a better condition close to normal. The average thicknesses of the choroid layer in DM rats were 15.69 ± 0.63, 14.87 ± 0.53, and 16.44 ± 0.57 µm, respectively (Fig 2). The mean wall thicknesses of choroidal arteries in the DM+GB group, DM+GM group, and DM+GR group were 10.23 ± 1.11, 10.41 ± 1.44, and 9.80 ± 1.78 µm, respectively. The mean diameters of the lumen were 52.52 ± 11.10, 53.71 ± 7.12, and 51.58 ± 8.11 µm, respectively, as shown in Table 4.

### The immunofluorescence method was utilized to investigate the VEGF and CD31 concentrations

During the research that was conducted on the expression of VEGF protein in the choroidal layer of the eye, it was discovered that the expression of VEGF protein in the choroid of the diabetic group rose in comparison with the control group, particularly in the choriocapillaris and the retinal pigment epithelium (RPE) (Fig 3, 5). It was found that the level of VEGF protein in the choroid layer of DM rats was statistically significantly higher than that of control rats (Fig 3). The mean fluorescence intensity of VEGF protein in the choroid of diabetic (DM) rats was 13.62+0.71, while in the control group (C), the mean VEGF protein fluorescence intensity was only 6.12+0.51. When studying the expression of VEGF protein in the choroid layer of rats in the DM+GB, DM+GM, and DM+GR groups (Fig 6A), it was found that the expression of VEGF protein was significantly decreased compared to rats in the diabetic group (DM), with the mean fluorescence intensity of VEGF protein being 7.96+0.77, 7.47+0.65, and 7.015+0.57, respectively.

**Fig 3.**
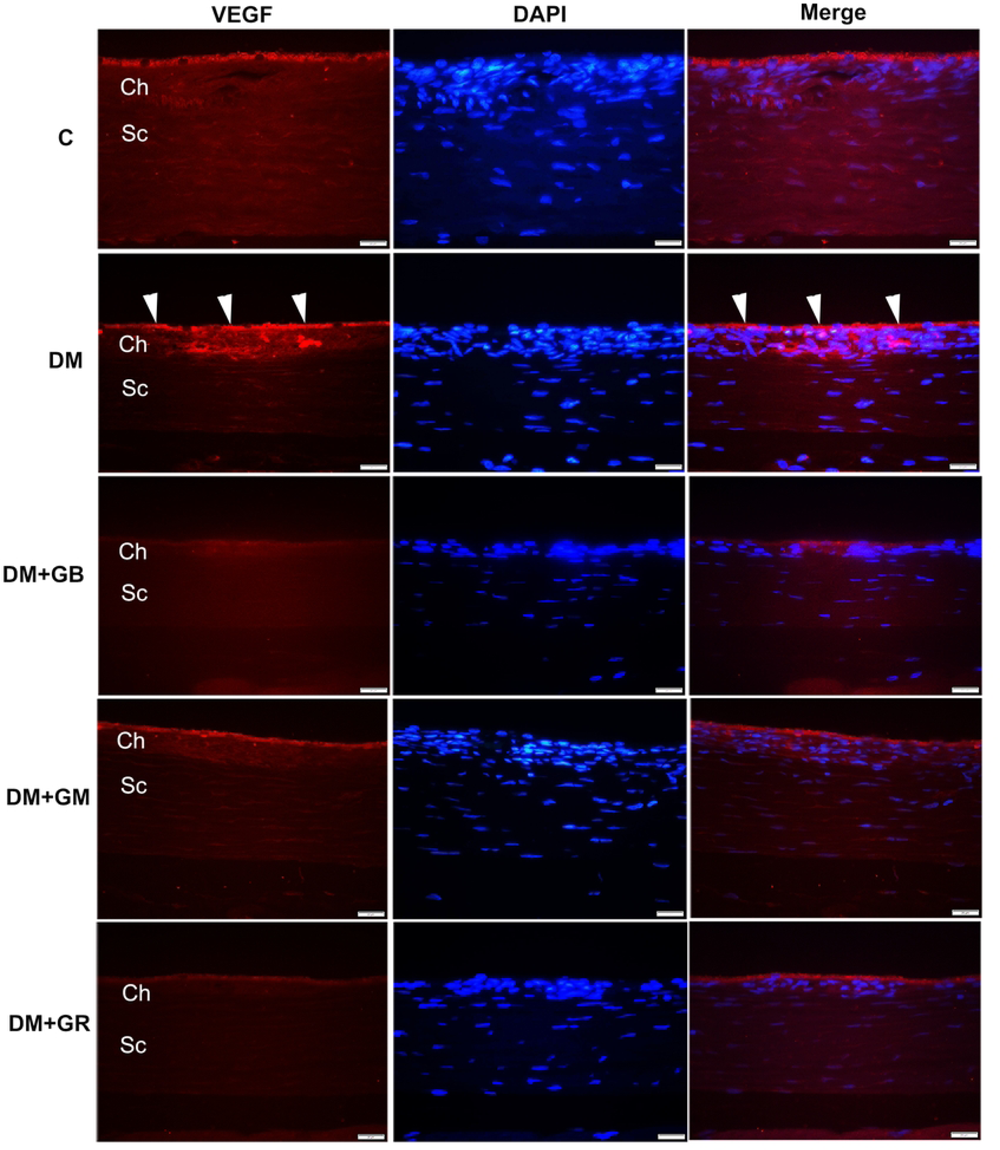
Glabridin and gymnemic acid can reduce a potent angiogenic cytokine. Representative images of the immunohistochemical sections of the choroidal layer with specific VEGF antibodies in rat eyes. Examples of immunoreactive VEGF, particularly in the choriocapillaris and the retinal pigment epithelium (RPE) (white arrowheads), were present in the DM group at 600 magnifications. Scale bar = 20 µm.

**Fig 4.**
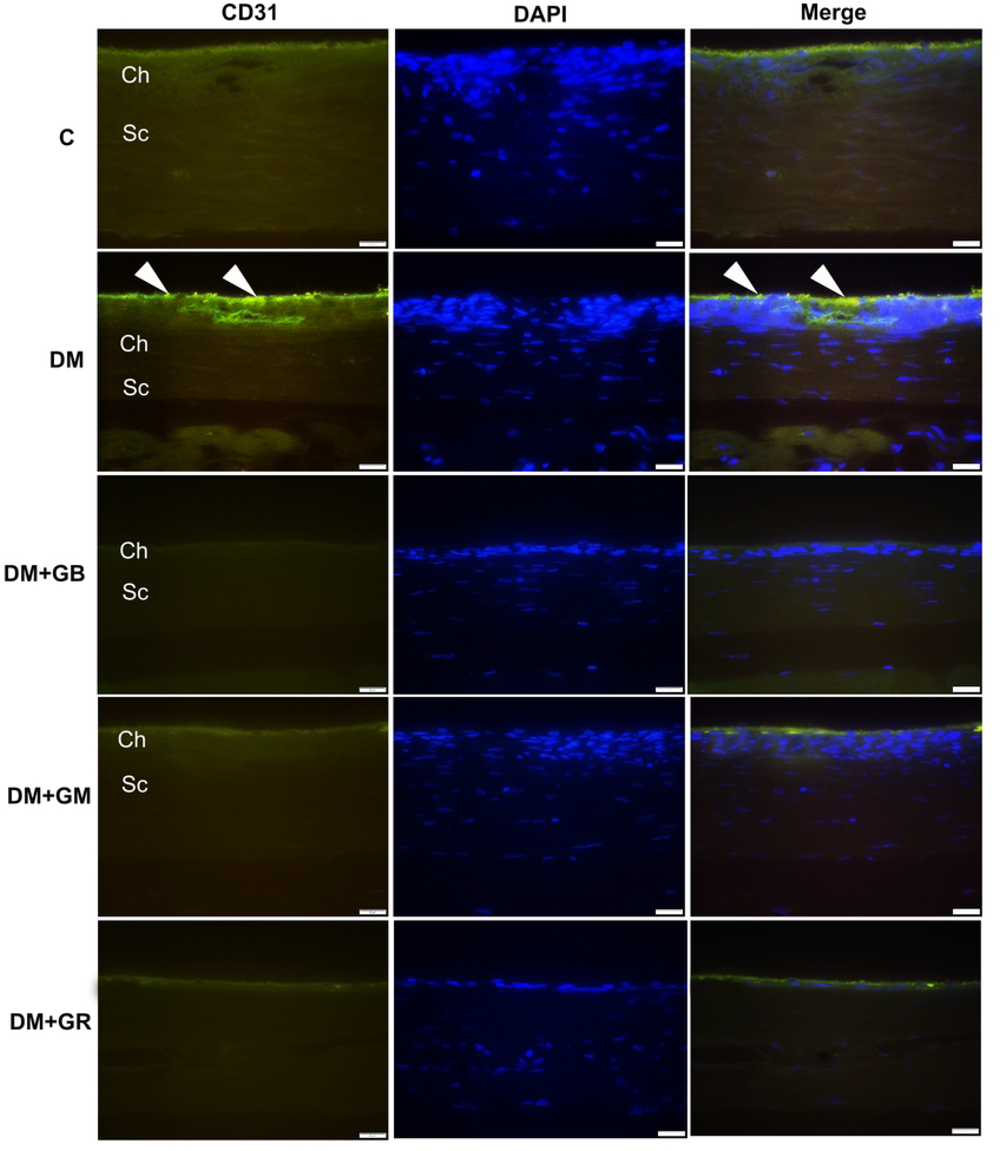
Glabridin and gymnemic acid have the ability to reduce the amount of angiogenic cytokine, which is essential for the process of survival. Representative images of the immunohistochemical sections of the choroidal layer with specific CD31 antibodies in rat eyes. Examples of immunoreactive CD31, particularly in the regions of the RPE and choriocapillaris as well as in the walls of blood vessels (white arrowheads), were present in the DM group at 600 magnifications. Scale bar = 20 µm.

**Fig 5.**
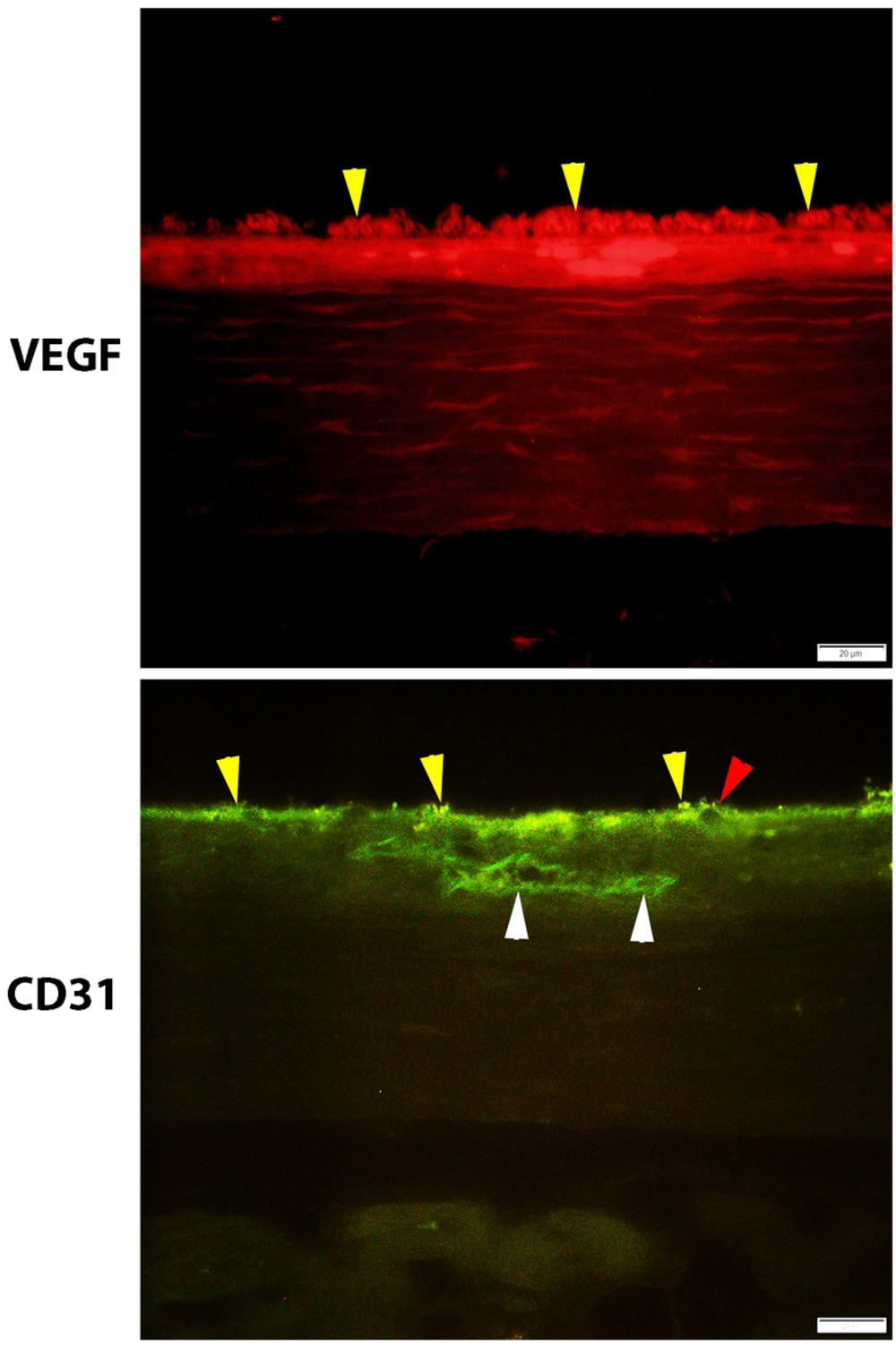
Photomicrographs of immunohistochemical sections of the choroidal layer with specific antibodies in the diabetic group. (A) An example of immunoreactive VEGF in the choriocapillaris and the retinal pigment epithelium (RPE) (yellow arrowheads). (B) Immunoreactive CD31 in the region of the RPE (yellow arrowheads), choriocapillaris (red arrowhead), and the wall of blood vessels (white arrowheads) was present in DM group. Scale bar = 20 µm.

An examination of the expression of CD31 protein in the choroid layer of diabetic rats revealed a statistically significant increase in the expression of CD31 protein (Figs 4, 5) in comparison to the expression of CD31 protein in control rats, particularly in the region of the RPE, choriocapillaris, and the wall of blood vessels. It was found that the mean CD31 protein intensity in the choroid layer of diabetic rats (DM) was 9.53+0.44, whereas the mean CD31 protein intensity in the control group (C) was 6.64+0.73 (Fig 6B). It was also found that the DM+GB, DM+GM, and DM+GR groups had lower levels of CD31 protein expression in the choroid layer compared to the diabetic (DM) group. The CD31 protein had a mean fluorescence intensity of 7.67+0.91, 7.99+0.65, and 8.66+ 0.94, respectively, from the three different samples (Fig 6B).

**Fig 6.**
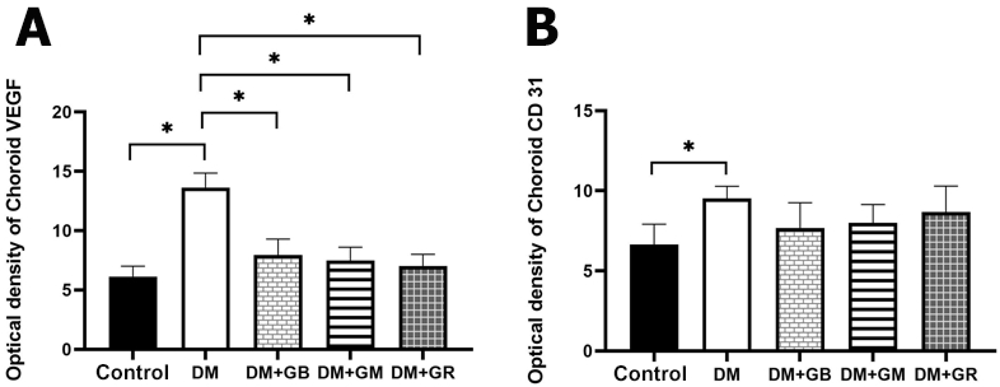
(A) The effect of glabridin and gymnemic acid on the Immunoreactive cytokines of the choroidal layer in five groups of rats at 8 weeks. A representative graph presents the relative VEGF optical density in the choroidal layer. Values are mean + SE, *p <0.001 and ** p <0.01, in the DM group when compared with the control, DM+GB, DM+GM, and STZ+GR groups. (B) The relative CD31 optical density in choroidal layer was analyzed. Values are mean + SE, *p <0.01, in the DM group when compared with the control group.

### Using vascular corrosion casting / scanning electron microscope to examine the anatomy of the choroidal blood vessels

In the choroidal walls of the eye, there is a posterior ciliary artery that connects to the choroid artery and divides into a network of capillaries known as the choriocapillaris. This network helps to transport blood throughout the eye. The choriocapillaris connects to the retina through the retinal pigment epithelium (RPE) and certain photoreceptor cells. Large-caliber vessels make up the outermost layer of the choroid, which is also referred to as Haller’s layer. Sattler’s layer is the name given to the innermost layer of the choroid (Fig 7), which is made up of vessels that are much smaller in size. Many anastomotic capillaries are what make up the choriocapillaris of the choroid, which is the part of the choroid that runs the deepest within the chest. For venous drainage to occur from the choriocapillaris, the vortex veins are the principal conduits that are utilized. Upon examination, it was discovered that the choroidal artery had branched out into the choriocapillaris in an organized fashion. In control rats, they were stacked in a thick manner, and there was no evidence of blood vessel fracture (Fig 8a). The average diameters of the choroidal artery and choriocapillaris in control rats were 89.73±2.82 and 10.52±0.72 µm, respectively (Table 5). In addition to being clearly visible under sparse vessels, the choriocapillaris alterations were more noticeable in DM rats. It was elongated and decreased in quantity, and it was also plainly visible. In the blood vessels that were destroyed in the DM group, choroidal arteries had a small diameter, stenosis was observed, and choriocapillaris shrank (Fig 8b). The arrangement of choriocapillaris was not dense or untidy. There was a connection in the choriocapillaris that was wilted and appeared as thin fringes with tube breaks. Within a particular region, the choriocapillaris had ruptured and linked to one another, resulting in the formation of a flatter sheet (Fig 9a). In addition, the sprouting of blood vessels was more identifiable in the choroidal arteries in Sattler’s layer (Fig 9b). The average diameters of the choroidal artery and choriocapillaris in DM rats were 60.31±2.71 and 4.68±0.65 µm, respectively, as shown in Table 5.

**Fig 7.**
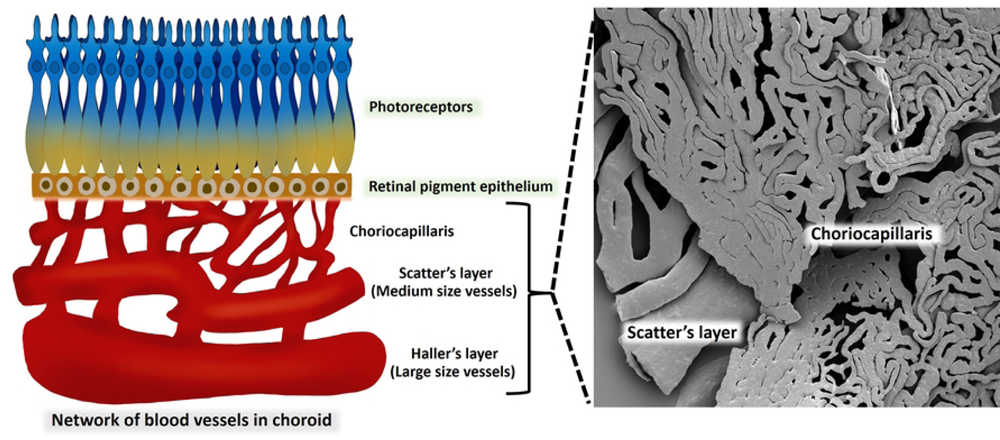
A comparison of the normal choroidal vascular structure and the vascular corrosion casting framework is indicated in this diagram. The outermost layer of the choroid, which is also known as Haller’s layer, is composed of vessels with a large vessel capacity. Sattler’s layer is the name given to the innermost layer of the choroid, which is composed of vessels that are significantly smaller in size than the vessels that make up the outermost layer. An extensive number of anastomotic capillaries are what constitute the choriocapillaris of the choroid.

**Fig 8.**
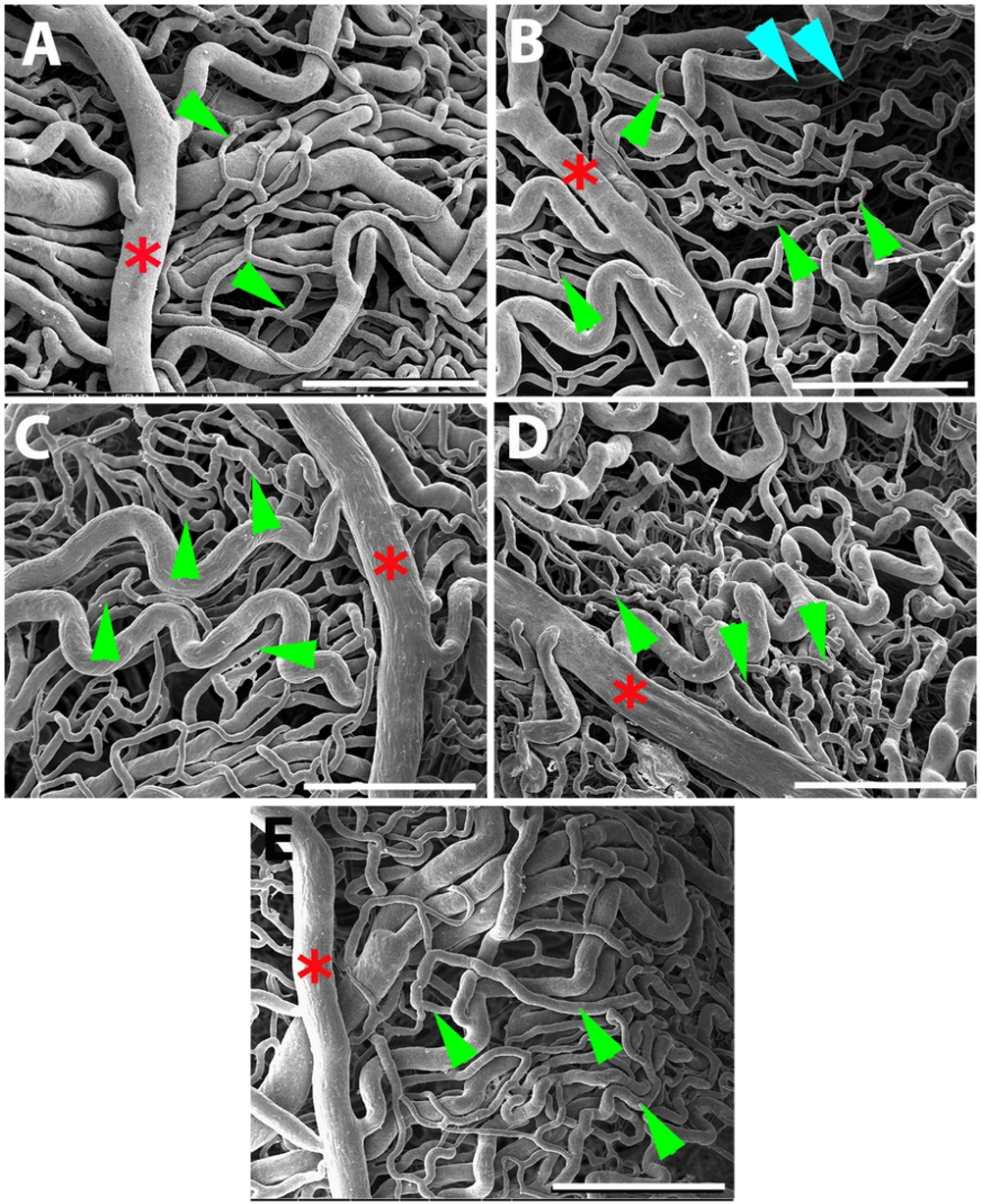
Glabridin and gymnemic acid have the capacity to enhance the choroid microvasculature feature. SEM micrographs with vascular corrosion casting technique/SEM in all five groups of rats: control (A), DM (B), DM+GB (C), DM+GM (D), and DM+GR (E). The choroid artery (red asterisk) splits into branches to supply the choroidal layer. Many anastomotic capillaries are what make up the choriocapillaris of the choroid (green arrows). In the DM group, choroid artery had a small diameter and the choriocapillaris shrank. The capillaries dropping out of the choriocapillaris (blue arrowheads) were also present in DM group. Scale bar = 300.

**Fig 9.**
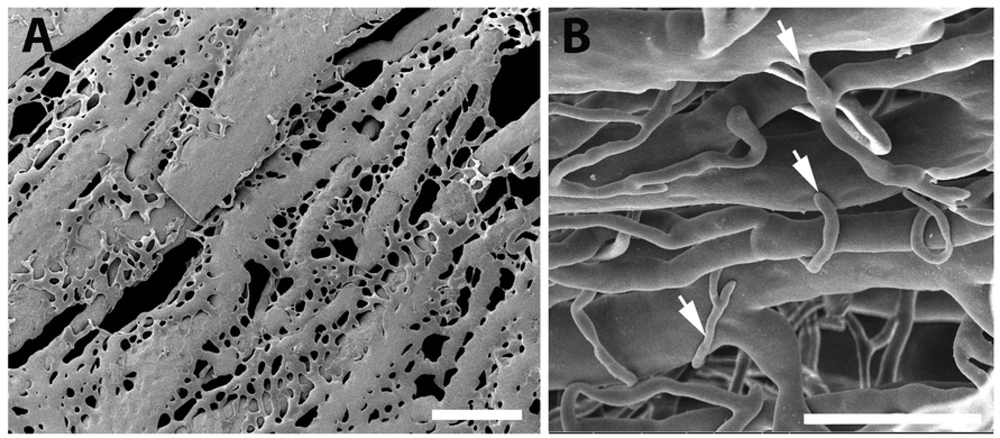
SEM micrographs using a vascular corrosion casting technique/SEM in the DM group. (A) The arrangement of the choriocapillaris is not dense and untidy; they link to one another, resulting in the formation of a flat sheet. (B) Furthermore, the sprouting of blood vessels (white arrows) was discovered in Sattler’s layer, which included choroidal arteries that were significantly more exposed. Scale bar = 300.

**Table 5.** The average diameters of choroid arteries (CRA) and choriocapillaris in the control, DM, DM+GB, DM+GM, and DM+GR groups at 8 weeks.

| Groups | Diameters of blood vessels ($\mu\text{m}$), mean ± SE | |
| --- | --- | --- |
|  | Choroidal arteries | Choriocapillaris |
| Control | $89.73 \pm 2.82$ | $10.52 \pm 0.72$ |
| DM | $60.31 \pm 2.71^*$ | $4.68 \pm 0.65^*$ |
| DM+GB | $81.18 \pm 3.56$ | $8.88 \pm 0.70$ |
| DM+GM | $80.31 \pm 3.83$ | $7.76 \pm 0.82$ |
| DM+GR | $79.82 \pm 4.73$ | $9.54 \pm 0.78$ |
| P-value* | $< 0.001$ | $< 0.001$ |

SE = standard error of mean.

In contrast, it was found that rats in the DM+GB group, DM+GM group, and DM+GR group were exposed to glabridin, gymnemic acid, and glyburide, respectively. It was found that blood vessels increased in diameter and were close to normal. The average diameters of the choroidal artery and choriocapillaris in the DM+GB group were 81.18±3.56 and 8.88±0.70 µm, respectively. The average diameters of the choroidal artery and choriocapillaris in the DM+GM group were 80.31±3.83 and 7.76±0.82 µm, respectively. The average diameters of the choroidal artery and choriocapillaris in the DM+GR group were 79.82±4.73 and 9.54±0.78 µm, respectively, as shown in Table 5.

## Discussion

From the results of this study, after the end of the 8-week experiment, rats in the diabetes group (DM) had higher blood sugar levels and significantly de-creased body weight. Compared to the control group (C), consistent with the results of a 2021 study by Sandech et al., the effects of blood sugar levels and body weight were studied. Streptozotocin at a dose of 60 mg/kg BW induces diabetes in rats. The experiment was conducted for a period of 8 weeks. It was found that the blood sugar level of the diabetic rats increased, and the body weight of the rats decreased with statistical significance compared to normal rats due to streptozotocin (10). Because of its low cost and the fact that it has less adverse effects than other medications, it is one of the diabetogenic medications. Specifically, the process of the alkylation of DNA is linked to the specific harmful impact that it has on β-cells that are located within the pancreas (14). Because of this, the production of the hormone insulin is slowed, which is a consequence of the action. This is accomplished via binding with GLUT2 to distribute streptozotocin, which can enter cells. The DNA strand is then modified by the addition of a methyl group, which leads to the destruction of beta cells and necrosis. The result causes certain conditions in the body, namely hypoinsulinemia and hyperglycemia, and as a result, the body has higher blood sugar levels (15). One of the obvious specific characteristics of diabetes is a decrease in body weight. As a result of the loss or deterioration of proteins in the body, the body fluid of diabetic rats decreased (16).

Results from this study show that after the end of the 8-week experiment, diabetic rats treated with glabridin at a dose of 40 mg/kg (DM+GB) showed a statistically significant decrease in blood sugar levels when compared to the rats in the diabetic group. The administration of glabridin at a dose of 40 mg/kg BW to rats induced with diabetes by injecting streptozotocin at a dose of 60 mg/kg BW was able to increase levels of antioxidants (superoxide dismutase; SOD) and reduce levels of free radicals (malondialdehyde; MDA), which causes the body to reduce oxidative stress, thus helping to reduce blood sugar levels (11, 12). In addition, this study shows that at the end of the 8-week experiment, diabetic rats treated with gymnemic acid at a dose of 400 mg/kg BW had a reduction in blood sugar levels compared to control rats. A study involving the administration of gymnemic acid at 400 mg/kg BW to rats induced with diabetes by streptozotocin injection at a dose of 60 mg/kg BW for a period of 60 days found that the blood sugar levels of diabetic rats were significantly reduced compared to untreated diabetic rats (10). The structure of the gymnemic acid molecule is somewhat comparable to that of the glucose molecule. There-fore, it can bind to the receptor instead of the sugar molecule in the taste buds on the tongue, inhibiting the taste of sweetness. It also binds to receptors in the small intestine and can delay the absorption of glucose into the small intestine (9). In addition, the administration of gymnemic acid can increase the number of beta cells in the pancreas. This is the reason that gymnemic acid can reduce blood sugar levels.

In the results of a study comparing the histological changes of cells and choroidal tissues in the eyeball using H&E and Masson’s trichome staining, it was found that the thickness of the choroid layer in the diabetic (DM) rats was significantly increased when compared to the control rats. This is consistent with research published in 2020 by Endo H. and colleagues who studied the thickness of the choroid layer in diabetic patients. The thickening of the choroidal layer is caused by the increased permeability of the choriocapillaris (17). Considering that the choroid is responsible for the blood flow to the eye, the autonomic nervous system plays a significant role in the autoregulation of the choroidal blood flow. During the early stages of DR, sympathetic innervation was engaged, which resulted in increased choroidal circulation, which ultimately led to an increase in the thickness of the choroid. At the same time, blood vessels in the choroid layer caused a significantly increased thickness of the blood vessel wall, especially in the connective tis-sue layer or the tunica adventitia, when compared to the control group rats. Hyperglycemia is directly responsible for the acceleration of the atherosclerotic process, which in turn leads to the development of endothelial dysfunction. This dysfunction, in turn, leads to vasoconstriction, proinflammatory processes, and prothrombotic processes, all of which contribute to the development and rupture of blood vessels (18).

Diabetes may be an independent factor that contributes to the thickening of the choroid, and subsequent progression of diabetic retinopathy (DR) may result in the reduction in choroid thickening. This may manifest as a thicker choroidal layer during the first stage of diabetic retinopathy (DR), which then gradually thins out as the disease progresses (19). The increased permeability of the choriocapillaris is what causes the thickening of the choroidal layer. Angiogenesis and cytokines that are caused by inflammation and oxidative stress may be the cause of the thickening of the choroidal layer in the early stages of diabetic retinopathy. This is because there is a possibility that these cytokines produce an excessive quantity of expression. These cytokines include platelet-derived growth factor, monocyte chemotactic protein-1, VEGF, pigment epithelium-derived factor, and insulin-like growth factor 1, among others (17). Using Masson’s trichrome staining to examine the choroidal arteries from the choroidal tissues of diabetic rats revealed that the vessel wall had become thicker and there were more collagen fibers deposited in the wall of the vessel. A narrowing of the lumen was also observed in rats with diabetes. The buildup of collagen can result in the stenosis of the arteries. Recent collagen synthesis has the potential to act as a substrate, which can result in luminal narrowing (20). A narrowing of the arteries that provide blood to the body, particularly the head, face, and brain, is referred to as arterial stenosis (21). In diabetic patients, vascular problems are the result of the deterioration of the vascular wall (21). Since these histopathological alterations cause blood vessels to become rigid, they are unable to respond to either exogenous or endogenous stimuli, which prevents them from being able to efficiently regulate blood flow (22).

This study used induced diabetic rats as subjects and used SEM to show that the choroidal arteries were changed in a way that was not seen in any of the other five groups of rats. Choroidal damage is increasing in rats with diabetes mellitus. The choroid micro-vasculature is made up of choroid arteries, choriocapillaris, short-running arterioles and venules, and choroid arteries. The choroid arteries and the choriocapillaris were observed to have tortuosity, shrinkage, and widespread constriction. The choriocapillaris, which is the smallest vessel, was the most severely affected by the damage. In diabetic eyes, the choriocapillaris can become blocked, the blood vessels can change shape with more vascular tortuosity, the blood vessels can drop out, there can be areas of vascular non-perfusion, and the choroidal neovascularization can occur. These findings were strikingly comparable to those observed in a previous study. Hidayat and Fine (23), who were the first to propose the idea of diabetic choroidopathy, discovered capillary dropout and choroidal neovascularization (CNV) in the enucleated eyes of diabetic patients. They used light and electron microscopy to make their observations. When the choriocapillaris is damaged, it can cause substantial damage to the function of the retinal tissue, particularly in the macula fovea.

All the gaps between the capillaries were not regular. Both the arterioles and the venules exhibited a significant amount of loss. Given the observations, there is indication that choroidal neovascularization has taken place. Higher blood glucose levels are the main cause of microvascular complications in diabetes, such as retinopathy, nephropathy, and neuropathy. Numerous studies have demonstrated that hyperglycemia has a detrimental effect on the endothelium’s functioning and causes pathological alterations that are associated with diabetes. In endothelial cells, hyperglycemia activates four primary molecular signaling systems. These mechanisms are in the cell membrane. Some of these turn on PKC, speeding up the hexosamine pathway, making more advanced glycation end-products, and speeding up the polyol pathway (24). PKC, for instance, affects the activation of a variety of growth factors and alters the activity of vasoactive factors in the context of diabetes microvascular problems. Vasoactive factors include vasodilators like nitric oxide (NO) as well as vasoconstrictors like angiotensin II and endothelin 1. These vasoactive factors are responsible for relaxing blood vessels.

VEGF plays a significant role in the process of neural regeneration and angiogenesis that takes place after an ischemic stroke (25). In addition, it has been demonstrated that VEGF, which is the main regulator of angiogenesis, may directly modulate the size of lumens. Lumen diameter is a reaction to blood pressure and blood flow, which influence the delivery of oxygen and immune surveillance that can occur. Furthermore, VEGF and Ang I are capable of inducing hyperplasia, tortuosity, and a reduction in the lumen diameter. These effects are highly potent (10). They are also quite good at inducing endothelial proliferation. It is for this reason that the increasing quantity of these two proteins causes the diameter of the lumen to become more constricted. In addition, it has been demonstrated that VEGF, which is the main regulator of angiogenesis, may directly modulate the size of lumens.

The mechanism, in conjunction with the decreased permeability of Bruch’s membrane, which is observed with advancing age, causes a reduction in the amount of oxygen that is supplied to the retina as an individual ages (26). However, this can also result in choroidal neovascularization. Retinal and retinal pigment epithelial cells are responsible for the release of VEGF, which results in the dilation of the choroidal capillaries and an increase in blood flow. Diabetes, in general, is associated with decreased cellular proliferation and EC dysfunction, which causes angiogenesis to be impaired (27). The pathogenesis of DR has been linked to several pro-angiogenic cytokines, such as insulin-like growth factor I (IGF-1) and platelet-derived growth factor (PDGF). However, VEGF is generally acknowledged as the most important cytokine in the process of driving DR. Ren-in-angiotensin and peroxisome proliferator-activated receptor gamma are two more pathways that are taken into consideration. ROS levels rise, which creates IL-1, IL-6, IL-8, MCP-1, iNOS, IP-10, MMPs (especially MMP9), C5-9, and TNF-α. These ROS cause inflammation at the cellular level (28). ICAM-1 (CD54) or E-Selectin (CD62E); VCAM-1 (CD106); and PECAM (CD31) are examples of endothelial adhesion molecules that are in-creased in endothelial cells. CD31 is considered a potential target for atherosclerosis since it is considered a proinflammatory and, thus, a proatherosclerotic molecule. Additionally, CD31 is involved in a wide variety of other biological processes, such as angiogenesis, apoptosis, platelet aggregation, and thrombosis (29). It is important for the vascular endothelium to be able to survive and respond properly to different types of mechanical, immune, and metabolic stresses (30).

Additionally, the beneficial therapeutic impact of gymnemic acid, which is the ac-tive component generated by *Gymnema sylvestre*, has been examined, and it was discovered that gymnemic acid was present in every region of the plant. Rats that were given gymnemic acid had lower levels of reactive oxygen species (ROS) and higher levels of an-tioxidants like glutathione (GSH), glutathione peroxidase (GSH-Px), catalase (CAT), and malondialdehyde (MDA), and these levels decreased (31). The glycolysis pathway relies heavily on the enzyme known as gymnemic acid. It has been discovered that it interacts with glyceraldehyde-3-phosphate dehydrogenase. A favorable therapeutic action of glabridin isolated from licorice is that it increases the activity of superoxide dismutase (SOD) considerably, while simultaneously lowering the levels of malondialdehyde (MDA) in the liver, kidney, and pancreas. The compound known as glabridin is a powerful an-ti-inflammatory drug, antioxidant, and free radical scavenger (32). So, the hypoglycemic effects of glabridin and gymnemic acid in rats that were given STZ may be linked to the antioxidant effects of these two compounds, at least in part.

## Conclusions

This study demonstrates that diabetic rats with high blood sugar levels are more likely to exhibit pathological alterations in the choroid of the eye than rats without diabetes. It was found that when animals were given glabridin at a dose of 40 mg/kg and gymnemic acid at a dose of 400 mg/kg, the levels of VEGF and CD31 proteins decreased. This treatment also supported the recovery process. Glabridin and gymnemic acid are re-sponsible for the histological condition that affects the tissues and blood vessels in the choroidal layer. It has the impact of lowering the choroid thickness, reducing the thickness of the wall and lumen of choroid vessels, and lowering neovascularization, all of which are significant contributors to the development of diabetes in the eyes. The use of glabridin, an important extract from licorice, and gymnemic acid, an important extract from *Gymnema sylvestre*, to treat, ease, or prevent diabetic choroid disease is therefore something that should be implemented as it constitutes an additional therapy alternative. In addition, obtaining a wide range of information and expertise on this topic will be advantageous to patients in the future.

## Author Contributions

Conceptualization, W.K. and M.K.; methodology, W.K.; software, M.K.; validation, W.K., U.M., and M.K.; formal analysis, U.M. and M.K.; investigation, W.K. and U.M.; resources, U.M. and M.K.; data curation, M.K.; writing— original draft preparation, W.K.; writing, review and editing, W.K.; visualization, U.M.; supervision, W.K.; project administration, W.K.; funding acquisition, W.K. All authors have read and agreed to the published version of the manuscript.

## Informed Consent Statement

Not applicable.

## Data Availability Statement

The data presented in this study are available on request from the corresponding author.

